# The Defensive Behavior of Indiana Mite-Biting Honey Bees Against *Varroa destructor* and the Structure of Bee Mandibles

**DOI:** 10.1101/2020.12.07.403147

**Authors:** Jada Smith, Xaryn L. Cleare, Krispn Given, Hongmei Li-Byarlay

## Abstract

The honey bee (*Apis mellifera*) is the most important managed pollinator to sustainable agriculture and our ecosystem. Yet managed honey bee colonies in the United States experience 30–40% losses annually. Among the many biotic stressors, the parasitic mite *Varroa destructor* is considered one of the main causes of colony losses. Bees’ mite-biting behavior has been selected as a *Varroa*-tolerant or *Varroa*-resistant trait in the state of Indiana for more than a decade. A survey of damaged mites from the bottom of a colony can be used as an extended phenotype to evaluate the mite-biting behavior of a colony. In this study, on average, 37% of mites sampled from the breeding stocks of 59 colonies of mite biters in Indiana were damaged or mutilated, which is significantly higher than the 19% of damaged mites found in commercial colonies in the southern United States. Indiana mite biter colonies had a higher proportion of damaged mites, although among the damaged mites, the number of missing legs was not significantly higher. In addition, the morphology of pollen-forager worker bee mandibles was compared by X-ray microcomputed tomography for six parameters in two populations, and a difference was detected in one parameter. Our results provide scientific evidence that potentially explains the defensive mechanism against *Varroa* mites: structural changes in the worker bee mandibles.

## 1 Introduction

Since 1987, when the ectoparasitic mite *Varroa destructor* was first introduced in the United States, *Varroa* infestations have become the primary contributors to honey bee (*Apis mellifera*) colony losses (Guzmán-Novoa et al., 2010; Le Conte et al., 2010; Nazzi and Le Conte, 2016). Mature *Varroa* females are 1.1 × 1.5 mm in size, and males are 0.8 × 0.7 mm (Häußermann et al., 2015). Most of the mite’s life cycle happens inside the brood cells, including the egg, six-legged larva, protonymph, deutonymph, and adult developmental stages (Bailey, 1968; Genersch, 2010). *Varroa* mites can infest honey bee colonies and cause colony losses as they feed on the fat bodies of bee pupae and cause morphological and behavioral defects in bee development (Le Conte et al., 2010; Ramsey et al., 2019). Furthermore, the *Varroa* mite is an effective vector for the transmission of viruses within the honey bee colony (Di Prisco et al., 2011; Wilfert et al., 2016).

The European honey bee (*A. mellifera*) has developed a set of behavioral defenses against *Varroa* mites to keep the mite population low, such as grooming, biting, and performing hygienic behaviors (Arechavaleta-Velasco and Guzmán-Novoa, 2001; Guzman-Novoa et al., 2012; Ruttner and Hanel, 1992; Spivak, 1996; Tsuruda et al., 2012; Villa et al., 2017). The biting behavior of worker bees, which is also considered a type of grooming behavior, enables them to bite adult mites and remove the mites from their bodies (Peng et al., 1987; Pritchard, 2016; Ruttner and Hanel, 1992). Colonies selected for mite-biting behavior by instrumental insemination or open mating with feral colonies potentially will have greater fitness over subsequent generations. Field reports show that Indiana mite biters from Purdue University, which have been selected over the past decade, have a higher mean proportion of damaged mites in the breeding population compared with unselected Italian queen bee colonies from California (Andino and Hunt, 2011; Hunt et al., 2016; Morfin et al., 2019). However, no report to date has compared changes in the mandibles as a potential mechanism for the mite-biting behavior.

Breeding mite-resistant bees is critical to maintaining sustainable apiculture for local pollination and food and crop productivity (Oddie et al., 2017). The modern beekeeping and breeding technique of instrumental insemination enables honey bee queens and colonies to be artificially selected (Meixner et al., 2010). Breeding efforts have been made on different continents and in different countries to select for mite-tolerant or mite-resistant traits (Büchler et al., 2010; Rinderer et al., 2010; Spivak, 1996). Mite-resistant bees assist beekeepers in managing the growing chemical miticide-resistance problems, and they will play a critical role in promoting sustainable agricultural practices (Hamiduzzaman et al., 2017; Kanga et al., 2016). In the past, research breeding efforts were focused on different bee-breeding stocks to improve the colony health (Büchler et al., 2010; Guarna et al., 2015, 2017), including in Russian bees (Rinderer et al., 2010), *Varroa*-Sensitive Hygienic bees (Danka et al., 2011; Villa et al., 2017), and Minnesota Hygienic bees (Guarna et al., 2017; Spivak, 1996).

Honey bee mandibles are considered the main mouthpart that worker bees use to bite or chew parasites, including mites and wax moths in the colony (Ruttner and Hanel, 1992). Our chemical analysis revealed that 2-heptanone is secreted from the mandibles and that it acts as an anesthetic on wax moth larva and *Varroa* mites (Papachristoforou et al., 2012). Micro-X-ray-computed tomography (microCT) is a technology that enables fast three-dimensional (3-D) scanning in satisfactory spatial resolution without complicated and lengthy sample preparation procedures. In the past, this technique has been used to study the brain anatomy and evolution of bees, ants, and other insects (Coty et al., 2014; Larabee et al., 2017; Li et al., 2011; Ribi et al., 2008). However, microCT has not previously been used to determine the shape of the honey bee mandible in fine detail.

In this study, we hypothesized that the Indiana mite-biter breeding stocks would have a higher level of mite-biting behavior than commercial bee colonies. Our goal was to characterize the bees’ behavioral and morphological capacity for mite-biting behavior. The total number of mites, the percentage of damaged mites (as the parameter for determining mite-biting behavior), and the number of mite legs missing per colony were reported. In addition, the mechanism underlying the bees’ mite-biting behavior was investigated by examining the shape of the mandibles in 3-D and comparing Indiana mite-biter colonies with commercial colonies from the southern United States (mainly the state of Georgia).

## 2 Materials and Methods

### 2.1 Honey Bee Colonies

Fifteen colonies from five commercial sources were sampled from beekeepers who bought their package colonies in 2018, originally from the state of Georgia (from five different commercial providers). Mites were collected between September 19 and 26, 2018, from different areas in the state of Ohio (Figure 1, site a: one colony from Defiance, Ohio [Defianace1]; site b: 14 colonies from Bellbrook [NA1, NA2, NA3, NA5, NA6], Beavercreek [PBJohn1, PBJohn2], Cedarville [Dan4], and Wilberforce, Ohio [AB1, CSU23, CSU24, CSU32, CSU51, CSU52]). In total, 59 colonies of Indiana mite-biter honey bees were sampled on July 3, August 6, September 28, October 10, October 17, and November 9, 2018, at Lafayette, Indiana (Figure 1, site c; colony numbers are listed in Table S1 of the Supplemental Materials). Fresh mite samples were collected over a 5-day period from each colony at Purdue University’s main apiary. Some colonies were sampled twice. Seven colonies at the Wright Patterson Air Force Base (WPAFB), Huffman Prairie site housed seven virgin queens from Purdue stock colonies. All queens were open-mated with drones from feral colonies near WPAFB (within 2 miles). Mites from open-mated mite-biter colonies were collected from September 26 to 30, 2018.

**FIGURE 1.**
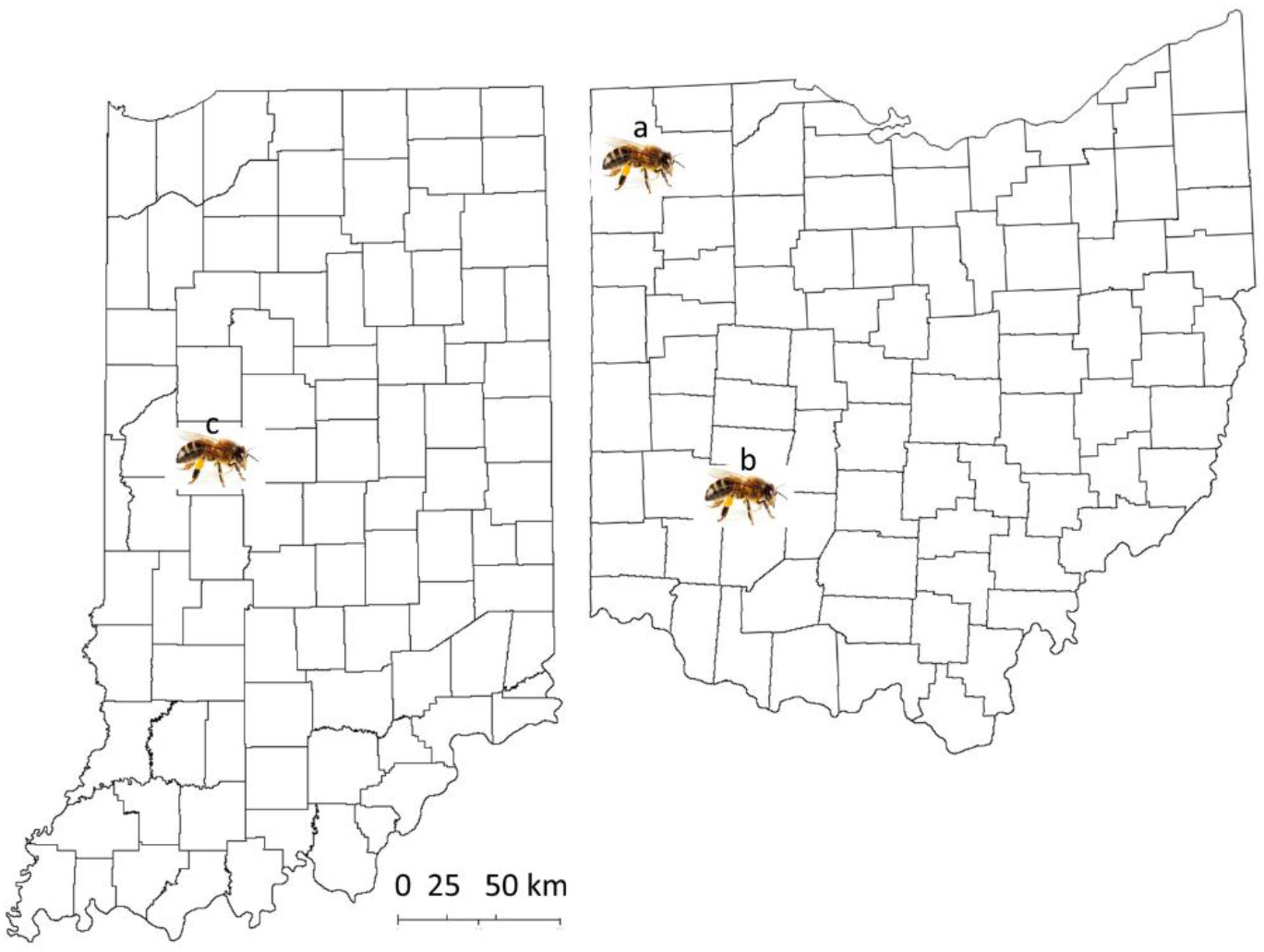
Map of the locations of sample collection from the states of Ohio and Indiana. Site a: Defiance, Ohio; site b: Green County, Ohio, including the cities of Bellbrook, Beavercreek, Cedarville, and Wilberforce; site c: West Lafayette, Indiana, for Indiana mite-biter colonies.

For all the worker bee samples used for mandible scanning, pollen foragers returning to the hive entrance (either a commercial or Indiana mite-biter colony) were collected with an insect vacuum (No. 2820GA, BioQuip Products, USA). All the bees were then frozen on dry ice, transported back to the laboratory, and kept in an –80 C freezer. At least 10 foragers per colony were collected from each site. Indiana mite-biter bee samples were collected in June 2018, and commercial bee samples were collected in July 2018.

### 2.2 *Varroa* Mites

Three groups of mites were compared: (1) commercial colonies, (2) mite-biter colonies (from Indiana), and (3) open-mated mite-biter colonies (at WPAFB). Figure 2 shows an example of a worker bee biting a *Varroa* mite on the top of a hive. Mite samples from Indiana mite-biter colonies were collected according to a previously described method (Andino and Hunt, 2011). For commercial bees and open-mated mite biters in Ohio, mite samples were collected as reported previously (Andino and Hunt, 2011). A small paintbrush was used to remove mites from the bottom boards into a plastic disposable cup with a lid (1 oz. volume). Mite samples stayed in a –20 C freezer overnight. Each mite was carefully glued onto a glass microscope slide (25 × 75 × 1 mm, Globe Scientific Inc.) with a small paintbrush. Slides were examined under a light microscope (Zeiss STEMI 580) with a magnification of 50×. Colonies with 15 or more mites sampled within a 5-day period were included in the data analysis. Mites collected in Ohio were examined for any missing legs (from 1 to 8) as visible damage in the viewer (Figure 3). The total number of mites sampled per colony and the number of damaged mites were compared among the three groups (commercial, Indiana mite biters, and open-mated mite biters). The number of missing legs was compared between commercial colonies and open-mated mite-biter colonies. Immature mites and empty mite body shells were excluded.

**FIGURE 2.**
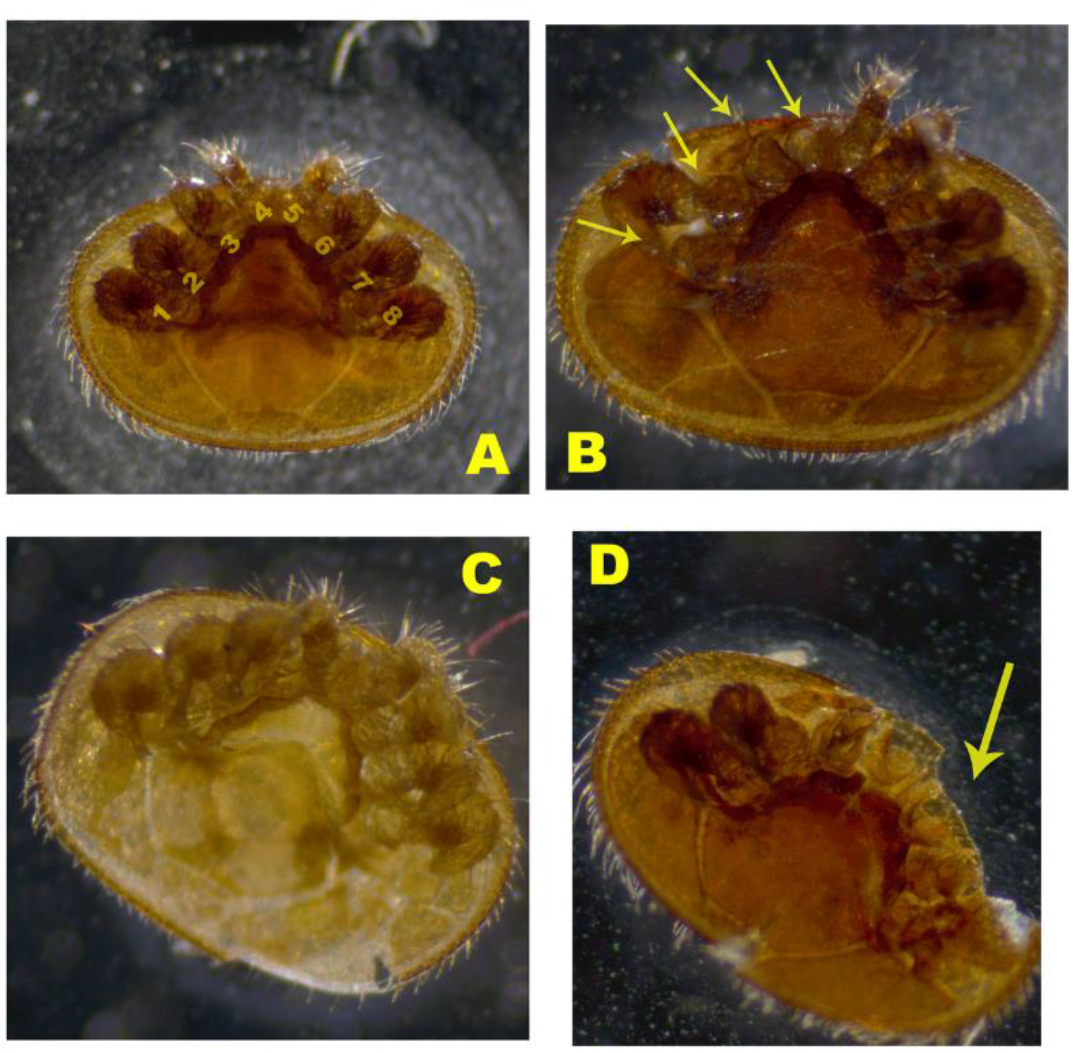
Images of damaged *Varroa* mites from the sampled colonies. **(A)** Mature mite with no damage. Label numbers 1–8 indicate the eight legs of the *Varroa* mite. **(B)** Damaged mite with legs missing. **(C)** Young mature mite. **(D)** Mature mite with a missing body part. Arrows indicate the damaged legs or body part.

**FIGURE 3.**
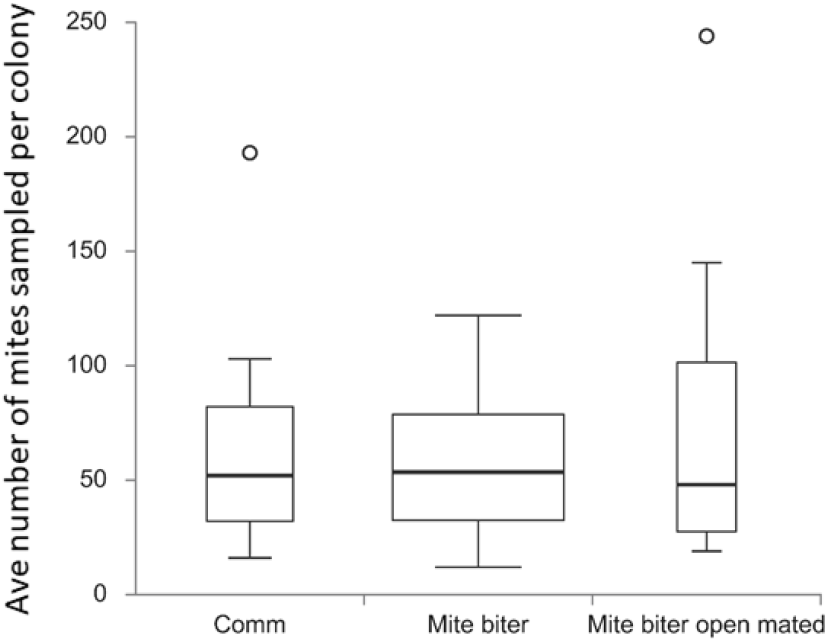
Box plots of the average number of mites collected per colony among the three groups: commercial colonies (Comm, *N*_colony_ = 15), Indiana mite-biter colonies (mite biter, *N*_colony_ = 59), and open-mated mite-biter colonies (open-mated mite biter, *N* = 7). The open circles are outliers of the colonies, *F*(2, 80) = 1.20, *p* = 0.31. The *p*-value corresponding to the *F*-statistic of the one-way ANOVA suggests that the three treatments were not significantly different.

### 2.3 MicroCT Scanning

From commercial colonies, five bees (five pairs of mandibles) were scanned from two colonies (three bees from colony PBJohn2 and two bees from colony Dan4). For the Indiana mite biters, nine bees from three colonies (three bees from colony 5, four bees from colony 15, and two bees from colony 41) were scanned. The scanning process was performed at the Center for Electron Microscopy and Analyses at Ohio State University (Columbus, Ohio) with a HeliScan microCT instrument (FEI Company, Thermo Fisher, USA) for 3-D imaging. A pair of mandibles was fixed to thin wooden posts (*r* = 1 mm, *h* = 148 mm) with superglue to fit in the HeliScan instrument. The scanning parameters were as follows: isotropic voxel size, 2.564 m per pixel; voltage, 60 or 80 kV; current, 80 or 46 µA; helical scan using space-filling trajectory reconstruction (Kingston et al., 2018) without any filter, 1,440 raw X-ray images. The software Avizo for FEI Systems (version 9.4, Thermo Fisher, USA) was used to quantify the measurements (height, length, width, small edge, long edge, and span of the spike area) for each scanned sample.

To compare the morphology of mandibles, six different parameters (Figure 4) were measured and compared between commercial colonies and mite-biter colonies. The height was measured from the top middle point to the base joint of the mandibular muscles. The long edge was the mandibular edge of the long side, similar to the blade on a pair of scissors. The short edge was the mandibular edge of the short side. The length was measured from the edge of the inner surface to the outer surface. The width was measured between the middle point of the long edge and the other side of the inner surface. The span of the spine area was the length of the sparse row of bristles or spines located along the inner side of the edge.

**FIGURE 4.**
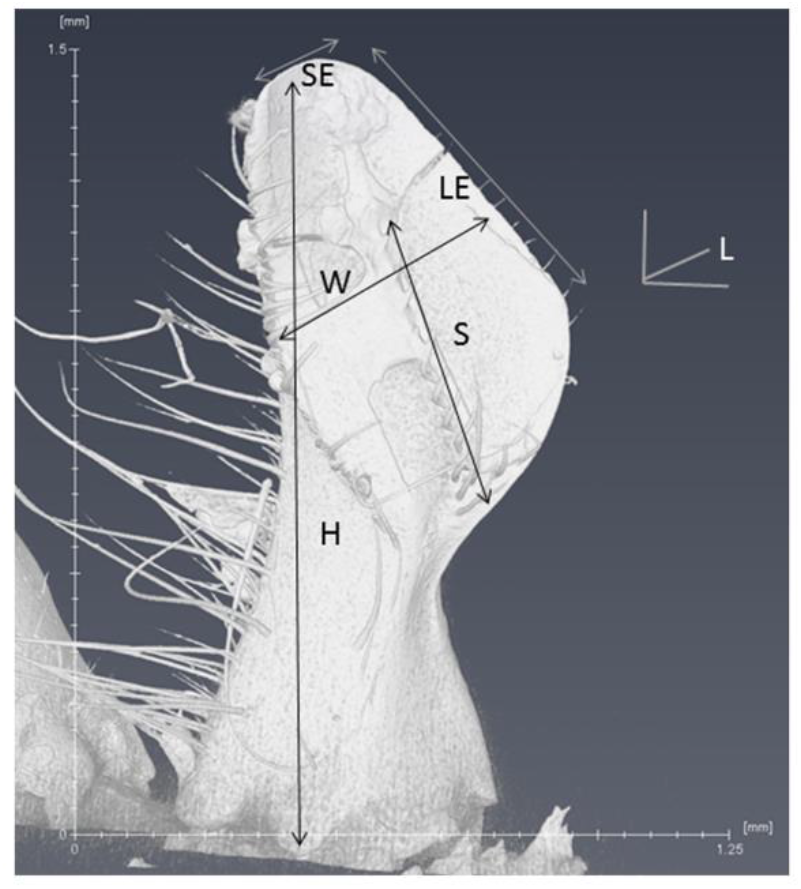
Violin plots of the percentage of damaged mites identified in relation to total mites collected per colony among three groups, commercial colonies (Comm, *N*_colony_ = 15, *N*_mite_ = 172), Indiana mite-biter colonies (mite biter, *N*_colony_ = 59, *N*_mite_ =1,201), and open-mated mite-biter colonies (open-mated mite biter, *N*_colony_ = 7, *N*_mite_ = 199). The one-way ANOVA suggested that one or more groups were significantly different, *F*(2, 80) = 28.86, *p* = 4.16e^−10^. The Tukey honestly significant difference test indicated that the levels of mite-biting behavior were not significantly different between the mite-biter and open-mated mite-biter colonies, but both were significantly higher than the commercial colonies, *Q* = 10.63 and 6.72, *p* < 0.01.

### 2.4 Data Analysis

The average number of mite samples per colony was compared among three groups, commercial bees, Indiana mite biters, and open-mated mite biters at WPAFB. If a colony was sampled twice in the fall season, the average number of mites collected was used for statistical analysis. The ratio of damaged mites to the total number of sampled mites was transformed by using arcsine [square root (*x*)] for normal distribution. A one-way analysis of variance (ANOVA), with post-hoc Tukey’s honestly significant difference (HSD) test calculated for comparing multiple treatments, was used to determine the differences among means of the different populations. Dependent variables were the total number of mites, the mite-biting rate, the number of legs damaged, and the mandible parameters. The online tool Interactive Dotplot (Weissgerber et al., 2017) was used to generate all the box plots.

## 3 Results

The mite-biting or grooming behavior referred to here involves a worker bee using its two forelegs and the two mandibles of its mouthpart to attack a *Varroa* mite in a colony. In addition to self-grooming, nest mates and groups of workers can actively remove adult female mites from worker bees and drop the damaged mites onto the bottom of the hive (Ruttner and Hanel, 1992). To characterize the damage to the mites, we categorized the observed mites into four different types: type A, mature adult female mites of a dark brown color with no damage and all eight legs present; type B, damaged adult female mites with legs missing; type C, young adult female mites of a pale color that were not counted as damaged adult mites; and type D, mites with body parts missing (Figure 2). Mite samples similar to type C were excluded because no damage was detected from worker bees’ grooming or biting behavior.

To evaluate the total mite population, we collected all the *Varroa* mites that appeared on the bottom board of a colony from each commercial colony (15 colonies, *N*_mite_ = 886), each mite-biter colony (59 colonies, *N*_mite_ = 3,390), and each open-mated mite-biter colony (7 colonies, *N*_mite_ = 569). No significant difference was detected among the commercial colonies, mite-biter colonies, and open-mated mite-biter colonies (one-way ANOVA of three independent treatments), *F*(2, 80) = 1.20, *p* = 0.31 (Figure 3).

To assess the bees’ mite-biting or grooming behavior, we surveyed all the damaged mites collected from the bottom board of each colony for all three groups of colonies (commercial colonies, *N*_mite_ = 172, mite-biter colonies, *N*_mite_ =1,201, open-mated mite-biter colonies, *N*_mite_ = 199). The means of the percentages of damaged mites per colony among the three colony types (commercial, mite biter, and open-mated mite biter) after transformation were 39.80%, 60.22%, and 60.14%. The one-way ANOVA suggested that one or more groups were significantly different, *F*(2, 80) = 28.86, *p* = 4.16e^−10^. The Tukey HSD test indicated that the levels of mite-biting behavior were not significantly different between the mite-biter and open-mated mite-biter colonies, but both were significantly higher than the level in commercial colonies, *Q* = 10.63 and 6.72, *p* < 0.01 (Figure 5). Commercial bees had the lowest mite-biting behavior among the three groups.

**FIGURE 5.**
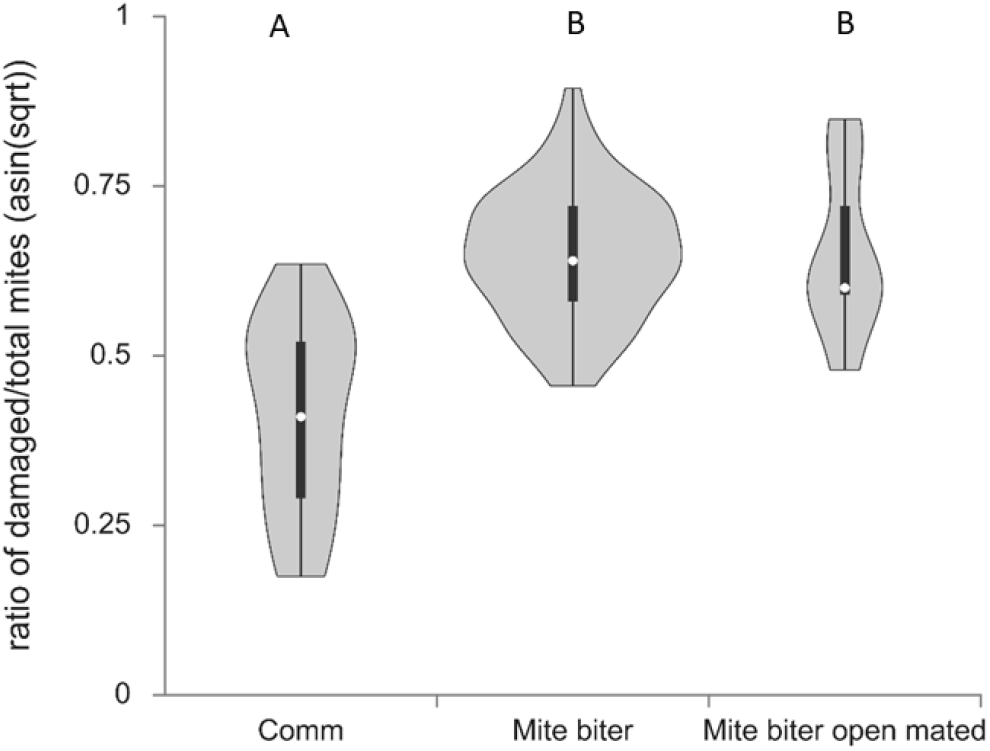
Box plots of the average number of legs missing per mite per colony among two groups, commercial colonies (Comm, *N*_colony_ = 15) and open-mated mite-biter colonies (open-mated mite biter, *N*_colony_ = 7). The open circles indicate outliers. The results showed no difference in the average number of legs missing per mite between the two groups, *Q* = 2.43, *p* > 0.05.

To further evaluate the bees’ damage to mites and the potential difference between commercial colonies and open-mated mite-biter colonies, we counted the number of legs missing from each damaged mite. This result showed no difference in the average number of legs missing per mite between these two groups, *Q* = 2.43, *p* > 0.05 (Figure 6).

**FIGURE 6.**
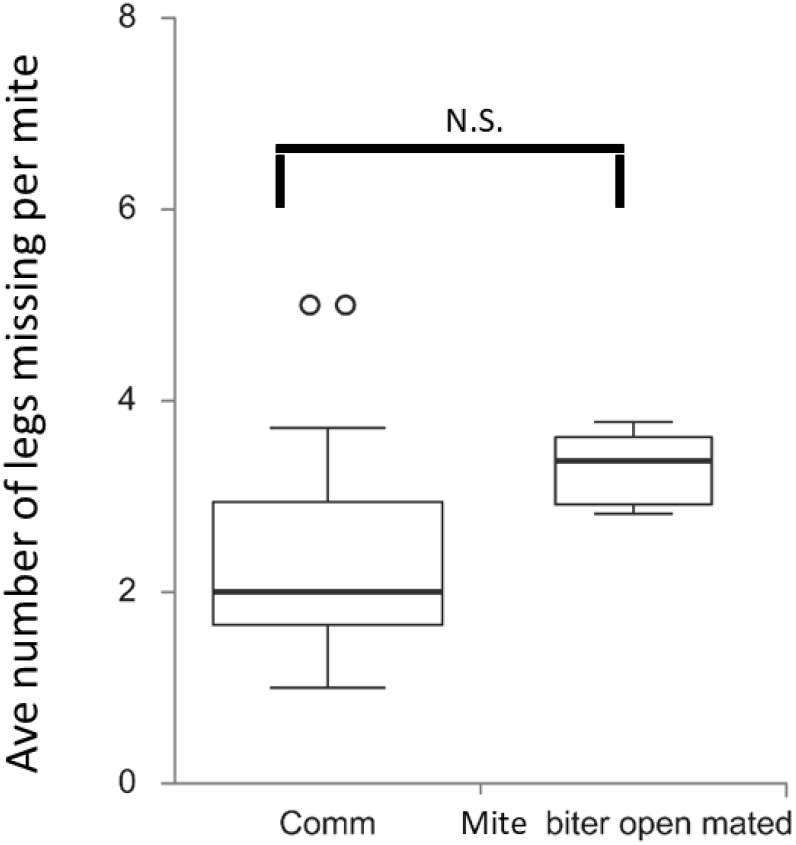
Six parameters of the scanned mandible are listed on the microCT image. L, length; W, width; H, height; SE, short edge; LE, long edge; S, span of the spine area.

To compare the morphology of mandibles between commercial colonies and mite-biter colonies, we measured six different parameters from the microCT data: the length, width, height, long edge, short edge, and span of the spine area (Figure 4). The ANOVA between these two groups showed that the long edge of mandibles in the mite-biter colonies were significantly shorter than those in commercial colonies, *F* = 5.78, *p* = 0.03 (Figure 7). We found no significant difference between the two groups in the other five parameters (Figure 7), but in the length, height, short edge, and span of the spine area, we noticed a consistent trend of smaller values in the mite-biter colonies.

**FIGURE 7.**
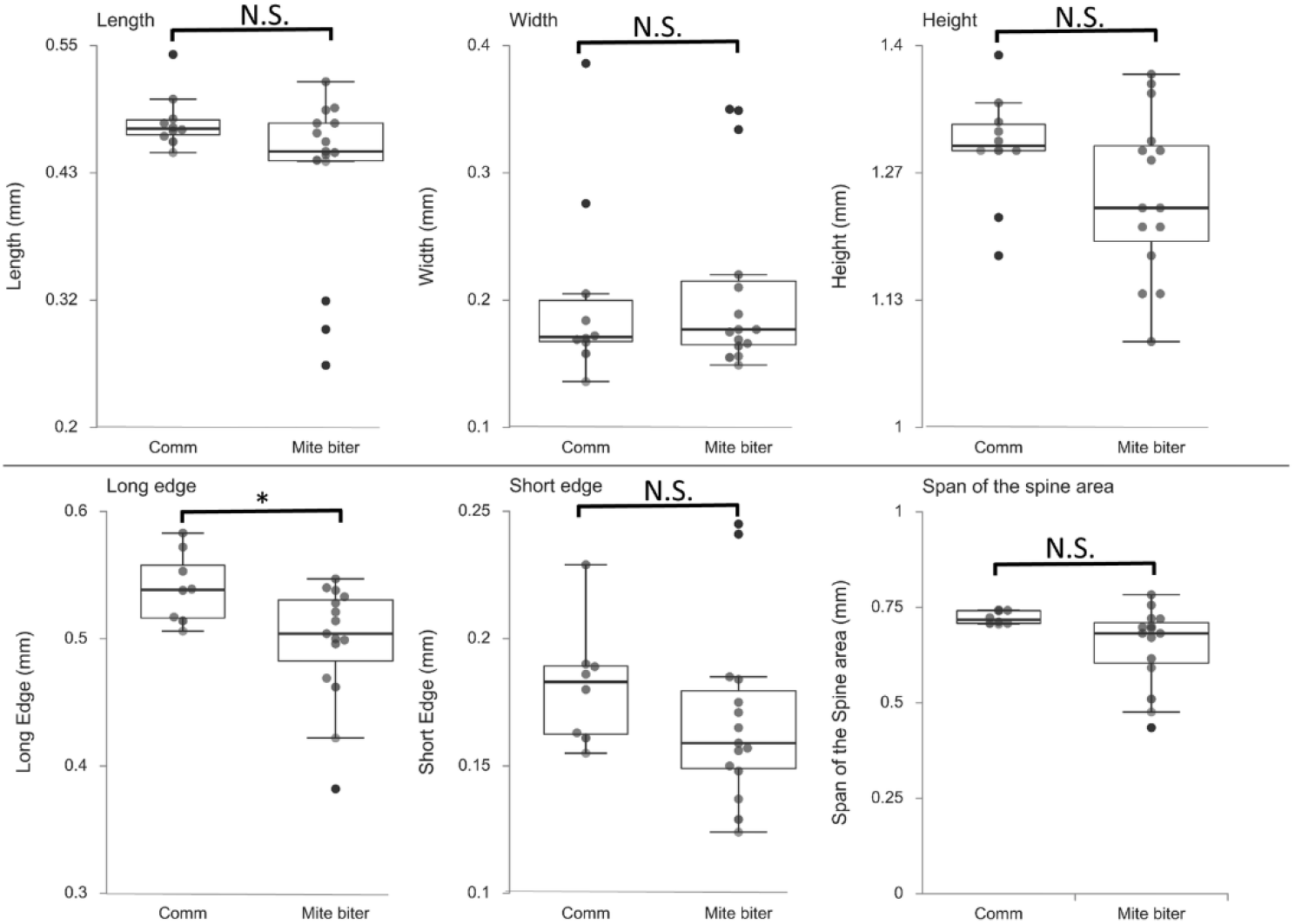
Box plots of the six parameters of mandible morphology between two groups, commercial colonies (Comm, *N*_mandible_ = 10) and mite-biter colonies (mite biter, *N*_mandible_ = 15). Each filled circle represents a data point. Results of the one-way ANOVA suggest that the two treatments were not significantly different in length, width, height, short edge, or span of the spine area, but they were significantly different in the long edge, *F*(1, 21) = 5.78, *p* < 0.05.

## 4 Discussion

We investigated the *Varroa* mite population and the differences in mite-biting behavior among commercial colonies, mite-biter colonies, and open-mated mite-biter colonies in the United States. In addition, we evaluated differences in the shape of bee mandibles between mite-biter stock colonies and commercial colonies. Bees in the mite-biter colonies displayed a higher level of mite-biting behavior than did those in the commercial colonies. The difference in the long edge of their mandibles may explain the physical mechanism by which their mandibles are able to mutilate mites.

Previous research on grooming behavior and damaged mites in *Apis* has shown that grooming behavior is a selected trait in naturally mite-resistant colonies (Arechavaleta-Velasco and Guzmán-Novoa, 2001; Boecking and Ritter, 1993; Fries et al., 1996; Peng et al., 1987; Russo et al., 2020). The mite-biting or grooming behavior of honey bees, as a defensive behavior against parasitic *Varroa* mites, can be used as a parameter to select for *Varroa* mite resistance in honey bee stocks (Hunt et al., 2016; Morfin et al., 2019; Pritchard, 2016; Rinderer et al., 2010; Spivak, 1996).

Our present comparison between commercial and mite-biter bees supports the value of selecting for mite-biting behavior in *A. mellifera*. With freshly damaged mites as our evidence, we provided a strong argument that workers of *A. mellifera* are able to amputate the legs of *Varroa* mites, as described by Ruttner and Hanel (1992). Collecting damaged mites from the bottom board of each colony within a 5-day time frame ensured that the observed damage on the mites was fresh. It is possible for beekeepers to record the proportion of damaged mites from the bottom boards of colonies and make a collaborative regional effort to select for the mite-biting trait in their region (Bienefeld et al., 1999; Hunt et al., 2016).

As a defensive response, honey bees use biting or grooming behavior to decrease the infestation of a mite population in the colony. Worker bees are known to use their mandibles to mutilate or damage *Varroa* mites, as reported previously (Ruttner and Hanel, 1992). Further studies have shown that certain chemicals, such as 2-heptanone, can be released from workers’ mandibles during a bite to anesthetize parasites in honey bee colonies, including *Varroa* mites and wax moths (Papachristoforou et al., 2012). Although bees’ mite-biting behavior has been reported, the underlying mechanism for this behavior has not been reported. Our results provide empirical evidence for changes in the structure of mandibles, such as the length of the long edge, which could be the mechanism underlying the biting behavior. In addition, our data showed that mite biters are under selection, which may lead to such structural changes toward mite resistance. Eastern honey bees (*A. cerana*) are the original hosts of *Varroa* mites, and in Asia, they have now evolved to be *Varroa* mite resistant. Their body size is also slightly smaller than that of *A. mellifera* (Peng et al., 1987; Yue et al., 2018). It is not clear if the change of body size is related to the evolution of biting behavior.

Our data indicate a clear trend that a behavioral adaptation is evolving in mite biters to defend against the parasitic mite *V. destructor*. These bees engaged in greater mite-biting behavior, perhaps because of their greater sensitivity to the mites, given the similarly high numbers of mite populations in all three colonies. Morfin et al. (2019) previously showed that the mite population was reduced in mite-biter colonies compared with colonies unselected for mite biting. However, our comparison of the total numbers of mites showed no significant difference among the commercial, mite-biter, and open-mated mite-biter colonies. This may be because the colonies we tested were in different geographic locations and had different management histories. Another possible mechanism for behavioral adaptation, via genetic changes such as the gene *AmNrx-1*, has been reported by Morfin et al. (2019).

The worker bees’ ability to detect mites may be based on their olfactory ability, considering that mite-biting behavior happens in dark hives most of the time. The mite-biter stocks may show a greater capability of detecting and recognizing mites as pests than do commercial colonies not selected for mite biting. Gradual changes may be taking place in the relationship between *A. mellifera* and *Varroa* mites. One potential change may be the biting mechanism of mite biters, one that Asian honey bees now display, as reported by Peng et al. (1987). Compared with the low frequency of mite removal and the limited success in clearing mites of bees in commercially sourced colonies, bees in mite-biter colonies exhibited an improvement in these abilities.

We identified the long edge of the mandible in bees from mite biter colonies as being shorter than that of bees from commercial colonies. The long edge is like a sharp knife that can be used to cut off the hind legs of *Varroa* mites. The pair of mandibles can act as a tool with double edges on the basal half of the rim. The mandibles adhere to the surface of mandibular muscles on the head. Potential differences of muscles may be related to the difference of biting ability among diffenert populations. These structures may explain why the change in the long edge affects the ability of workers to bite the mites. Even though other measurements did not show a significant difference, similar trends were observed in the short edge, height, and length.

Although evidence exists for variation in the mite-biting or grooming behavior of different genotypes (Guzman-Novoa et al., 2012), more research is needed on the genetic architecture and pattern of inheritance of this behavior for honey bee breeding and selection. With the rapid development of new sequencing technologies and genome editing tools, genome-wide marker-assisted selection may be applied in the future for honey bee breeding.

## Author Contributions

H.L.-B. conceived of and designed the experiments; J.S., X.L.C., K.G., and H.L.-B. collected samples and data. J.S. and H.L.-B. performed the data analysis and wrote the paper. All authors approved the final manuscript.

## Acknowledgments

We thank Greg Hunt, who guided sample collections at Purdue University and provided comments on the manuscript; Dwight Wells and Danielle Trevino for assistance in sample collections at the WPAFB; Nathan Arndt, Andy Bern, John Gazzerro, Danielle Kroh, and Jamie Walters for access to the commercial colonies; Carley Goodwin at Ohio State University for assistance with the microCT process; and Kaila Young at Central State University for assistant in mounting and scanning samples.

